# OT-RMSD: A Generalization of the Root-Mean-Square Deviation for Aligning Unequal-Length Protein Structures

**DOI:** 10.1101/2025.09.23.678064

**Authors:** Yue Hu, Zanxia Cao, Yingchao Liu

## Abstract

The Root-Mean-Square Deviation (RMSD) metric, coupled with the Kabsch algorithm for optimal superposition, represents a cornerstone of quantitative structural biology. However, its application is fundamentally limited to the comparison of structures with a pre-defined, one-to-one correspondence of equal length, precluding its direct use for the vast majority of biologically relevant comparisons involving proteins of different lengths or unknown equivalences. To overcome this limitation, we introduce OT-RMSD, a novel, parameter-light method that generalizes the RMSD for comparing unequal-length protein structures without a pre-defined alignment. Our approach recasts the correspondence problem as an Optimal Transport (OT) task, utilizing a fundamental, distance-based cost function to identify the most economical mapping between two protein structures. This transport plan is then used as a soft-weighting scheme within an iterative Sinkhorn-Kabsch framework to simultaneously solve for correspondence and optimal superposition. We test our method on a benchmark dataset of 40 challenging protein pairs and demonstrate its superiority over both the widely-used PyMOL ‘align’ command and a more complex, heuristic-driven variant of our own framework. On average, OT-RMSD identifies larger and more accurate alignment cores, achieving a significantly lower RMSD. We conclude that OT-RMSD provides a mathematically principled, robust, and effective extension of the classic RMSD concept, offering a powerful tool for modern structural analysis.

## 1 Introduction

The comparison of three-dimensional protein structures is a fundamental task in molecular biology, providing critical insights into evolutionary relationships, functional mechanisms, and drug design. For decades, the Root-Mean-Square Deviation (RMSD) has stood as the gold standard for quantifying the similarity between two superimposed structures. Its enduring prevalence stems from its simple, intuitive physical meaning: the average distance between corresponding atoms. The calculation of RMSD is inextricably linked to the Kabsch algorithm [1], an elegant closed-form solution for finding the optimal rotation and translation that minimizes the RMSD between two sets of corresponding points.

Together, the RMSD metric and the Kabsch algorithm form a powerful and ubiquitous toolkit. However, their utility is predicated on a critical prerequisite: a pre-defined, one-to-one correspondence between two structures of equal length. This assumption holds for specific tasks, such as comparing conformational changes in the same protein, but it breaks down for the more general and common problem of comparing two different proteins. Different proteins rarely have the same number of residues, and even when they do, the correct residue-level equivalence is unknown beforehand. This limitation has traditionally forced a decoupling of the alignment problem into two separate steps: first, a heuristic-based method (e.g., sequence alignment, dynamic programming, fragment matching) is used to establish a correspondence; second, the Kabsch algorithm is applied to this correspondence to calculate an RMSD.

This two-step process is fraught with challenges. The final RMSD is critically dependent on the quality of the initial, heuristically-determined alignment. A suboptimal set of correspondences will invariably lead to a misleading RMSD, regardless of the mathematical elegance of the subsequent superposition. This raises a fundamental question: can we unify the correspondence and superposition problems? Can we define a “Root-Mean-Square Deviation” when the correspondence is not given and the structures have different lengths?

In this paper, we propose a direct and principled answer to this question. We introduce OT-RMSD, a novel framework that generalizes the RMSD concept by recasting the correspondence problem within the mathematical framework of Optimal Transport (OT). Optimal Transport provides a powerful, parameter-light theory for finding the most efficient mapping between two distributions of mass. We model proteins as distributions of C-alpha atoms and seek the most economical “transport plan” that aligns one to the other.

Crucially, our core OT-RMSD method relies on the most fundamental cost function imaginable: the squared Euclidean distance between atoms. This choice avoids the complex, manually-tuned heuristics that underpin many existing alignment algorithms, resulting in a method that is both mathematically pure and highly effective. This transport plan, which represents a “soft” correspondence, is then used to weight the contribution of each atom pair in a generalized Kabsch algorithm. By iterating between solving the OT problem and the weighted superposition problem, our algorithm simultaneously converges to both the optimal correspondence and the optimal alignment.

The goal of this paper is to present the OT-RMSD method as a natural and powerful extension of the classic Kabsch algorithm. We will detail its theoretical underpinnings and demonstrate its practical effectiveness through a rigorous benchmark comparison against the widely-used PyMOL alignment tool and a more complex, heuristic-based variant of our own method. Our results show that the simplicity and mathematical rigor of OT-RMSD allow it to identify larger, more accurate alignment cores with lower RMSD values, establishing it as a valuable new tool for quantitative structural comparison.

## 2 Methods

### 2.1 The Classic RMSD and the Kabsch Algorithm

The classic approach to comparing two protein structures, *P* and *Q*, assumes two conditions: (1) they are represented by sets of C-alpha coordinates, *P* = *{p*_1_, *p*_2_, …, *p*_*N*_ *}* and *Q* = *{q*_1_, *q*_2_, …, *q*_*N*_ *}*, of equal size *N*; and (2) a one-to-one correspondence between points (*p*_*i*_, *q*_*i*_) is known. The goal is to find a rigid-body transformation (a rotation *R* and a translation *t*) that minimizes the RMSD, defined as:

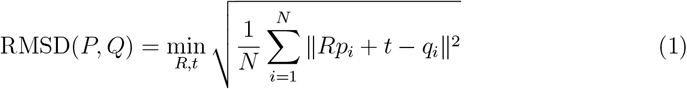

The Kabsch algorithm [1], also known as the Wahba’s problem in other fields, provides an elegant, closed-form solution to this minimization problem. Let’s break down the derivation step-by-step for clarity.

#### Step 1: Decoupling Rotation and Translation

The minimization problem involves both a rotation *R* and a translation *t*. We can simplify this by showing that the optimal translation *t* can be determined independently of the rotation. The term to minimize is the sum of squared errors, 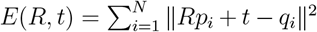. To find the optimal *t*, we can take the derivative of *E* with respect to *t* and set it to zero.

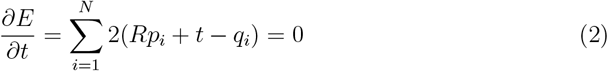

Solving for *t*, we get:

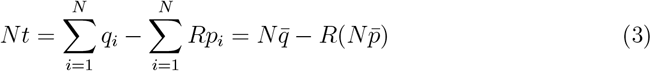

where 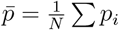 and 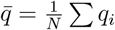 are the centroids of the point sets *P* and *Q*, respectively.

This gives us the optimal translation:

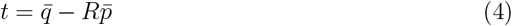

This shows that the optimal translation moves the centroid of the rotated structure *P* onto the centroid of structure *Q*. By substituting this back into the error function, the problem reduces to finding the optimal rotation *R* for the centroid-aligned structures. For simplicity, let’s assume from now on that both *P* and *Q* have been pre-centered by subtracting their respective centroids. The problem is now to minimize 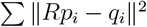.

#### Step 2: Formulating the Rotation Problem

We want to find the rotation *R* that minimizes 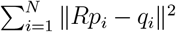. Let’s expand the squared norm:

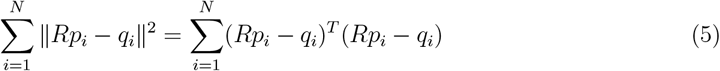

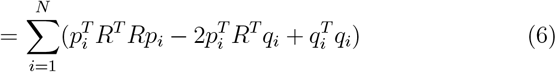

Since *R* is a rotation matrix, *R*^*T*^ *R* = *I* (the identity matrix). The expression simplifies to:

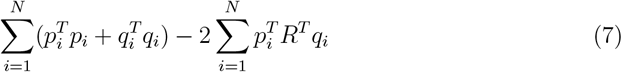

The first term, 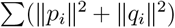, is a constant with respect to *R*. Therefore, minimizing. the entire expression is equivalent to maximizing the second term, 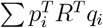trace trick, we can rewrite this as:

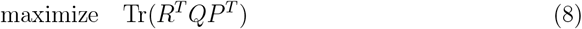

where *P* and *Q* are now *N* × 3 matrices whose rows are the coordinates 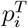 and 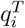.

#### Step 3: The Covariance Matrix and SVD

Let’s define the 3 × 3 covariance matrix *H* = *P*^*T*^ *Q*. The problem is to find the rotation *R* that maximizes Tr(*RH*). Now, we perform the Singular Value Decomposition (SVD) on the covariance matrix *H*:

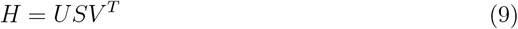

where *U* and *V* are 3*×*3 orthogonal matrices and *S* is a 3*×*3 diagonal matrix with non-negative singular values. Substituting this into our objective function:

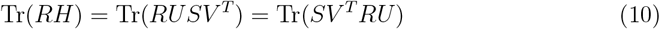

Let *Z* = *V* ^*T*^ *RU*. Since *V, R*, and *U* are all orthogonal matrices, their product *Z* is also an orthogonal matrix. Our goal is now to maximize Tr(*SZ*) subject to *Z* being orthogonal.

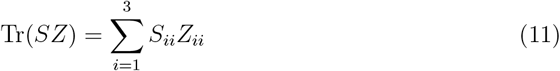

Since *S*_*ii*_ *≥* 0 and the elements of an orthogonal matrix *Z* are bounded by |*Z*_*ii*_| *≤* 1, this sum is maximized when *Z*_*ii*_ is as large as possible, which is *Z*_*ii*_ = 1. This occurs when *Z* is the identity matrix, *Z* = *I*.

#### Step 4: Finding the Optimal Rotation

If we set *Z* = *I*, we get *V* ^*T*^ *RU* = *I*. Solving for *R* gives:

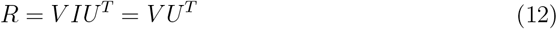

This is the optimal rotation matrix. However, we must ensure that *R* is a proper rotation matrix (i.e., det(*R*) = 1) and not a reflection (i.e., det(*R*) = *−*1). We can check the determinant of our solution: det(*R*) = det(*V U*^*T*^) = det(*V*) det(*U*^*T*^). If det(*V*) det(*U*) = 1, then our solution is a valid rotation. If det(*V*) det(*U*) = *−*1, it means our solution includes a reflection. In this “reflection” case, the optimal rotation is found by flipping the sign of the column of *V* corresponding to the smallest singular value in *S*. If we let *d* = sign(det(*V U*^*T*^)), the corrected rotation is:

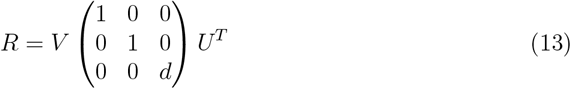

This correction ensures we find the best rotation in *SO*(3). With the optimal rotation *R* found, the optimal translation *t* is simply calculated as 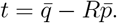

The limitation of this elegant algorithm is self-evident: the entire procedure hinges on the pre-supplied one-to-one correspondence, which is generally unknown when comparing different proteins.

### 2.2 The OT-RMSD Framework: A Principled Generalization

To overcome this limitation, we reformulate the alignment problem to solve for correspondence and superposition simultaneously. Our framework, OT-RMSD, is an iterative procedure that alternates between these two sub-problems, generalizing the Kabsch algorithm to cases of unequal length and unknown correspondence.

#### 2.2.1 Correspondence as an Optimal Transport Problem

We treat the two proteins, represented by C-alpha coordinates *P ∈* ℝ^*N×*3^ and *Q ∈* ℝ^*M×*3^, as two discrete probability distributions. The core of the alignment problem is to find a mapping, or “transport plan”, **P** *∈* ℝ^*N×M*^ that describes how to optimally map the “mass” of points in *P* to the points in *Q*. The entry **P**_*ij*_ in this matrix represents the amount of mass from point *p*_*i*_ that is transported to point *q*_*j*_. This matrix, which we call the P-matrix, represents a “soft” or probabilistic correspondence between the two structures.

#### 2.2.2 A Fundamental Cost Function

To define an “optimal” transport plan, we must first define the cost of transporting mass between any two points. Many alignment algorithms rely on complex, heuristic-based scoring functions. The OT-RMSD method, in contrast, is founded on the most fundamental and parameter-free cost function: the squared Euclidean distance. The cost of matching atom *p*_*i*_ with atom *q*_*j*_ is simply 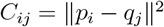.

This choice is central to our philosophy. By avoiding domain-specific heuristics, we create a method that is a pure and direct geometric extension of the RMSD concept. For the purpose of numerical stability and framing the problem as a maximization of rewards, we use a reward matrix *M* where the reward is inversely proportional to the cost:

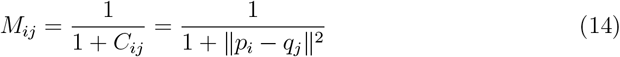

This formulation is simple, robust, and requires no free parameters.

#### 2.2.3 Iterative Solution via Sinkhorn-Kabsch

We solve for the alignment using an iterative algorithm that alternates between updating the transport plan **P** and the structural superposition (*R, t*).

### 1. Correspondence Update (Sinkhorn’s Algorithm)

Given the current superpo sition of structures, we calculate the reward matrix *M* as defined above. We then use the Sinkhorn-Knopp algorithm to find the optimal transport plan **P**. The Sinkhorn algorithm is an efficient iterative method for solving the entropy-regularized optimal transport problem. It takes the exponentiated reward matrix, *K* = exp(*γM*), where *γ* is a sharpness parameter, and iteratively normalizes its rows and columns until it converges to a doubly stochastic matrix. This resulting matrix is our transport plan **P**.

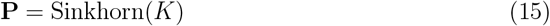

### 2. Superposition Update (Weighted Kabsch Algorithm)

With the transport plan **P** fixed, we now find the optimal superposition. The matrix **P** provides a “soft” correspondence, where **P**_*ij*_ acts as the weight for the atom pair (*p*_*i*_, *q*_*j*_). We use these weights in a weighted version of the Kabsch algorithm.

First, we compute the weighted centroids:

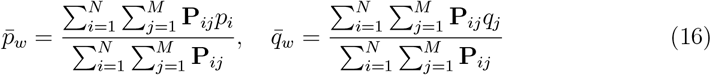

Then, we construct the weighted covariance matrix *H*_*w*_. This is the crucial step where the soft correspondence **P** is integrated. Let *P*′ be the *N ×* 3 matrix of centered mobile coordinates and *Q*′ be the *M ×* 3 matrix of centered target coordinates. The weighted covariance matrix is computed in a clean matrix form as:

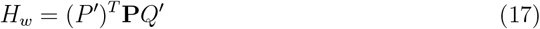

This is the key extension to the Kabsch algorithm. Instead of a simple covariance matrix between two sets of points with a one-to-one correspondence, we are now computing a covariance matrix that is weighted by the entire soft correspondence plan **P**. The SVD of this 3 × 3 matrix *H*_*w*_ is then used to find the optimal rotation, just as in the classic algorithm. The optimal rotation *R* is found using the same logic as in the classic Kabsch algorithm:

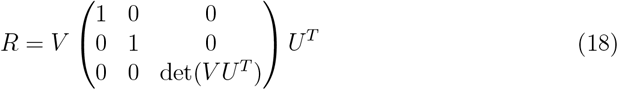

where *U* and *V* are the result of the SVD of *H*_*w*_ = *USV* ^*T*^. And the optimal translation is given by 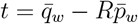.

### 3. Iterative Refinement

The overall OT-RMSD algorithm is a powerful iterative refinement scheme, akin to a coordinate ascent method. It alternates between optimizing two distinct sets of variables: the soft correspondence matrix **P** and the rigid-body transformation (*R, t*). In the first step of each iteration, we hold the geometric superposition fixed and solve the Optimal Transport problem to find the best correspondence plan **P**. In the second step, we hold this correspondence plan fixed and solve the weighted Kabsch problem to find the optimal superposition (*R, t*) that minimizes the RMSD under these soft correspondences.

Each of these steps is guaranteed to improve the overall alignment quality (specifically, to increase the total reward). By alternating between these two updates, the algorithm is guaranteed to converge to a self-consistent solution, where the correspondence and superposition are mutually optimal with respect to each other. This iterative process allows the algorithm to progressively and simultaneously discover both the shared structural core and the precise geometric transformation that aligns it.

#### 2.2.4 The OT-RMSD Metric and Core Identification

During the iterative process, we can monitor the convergence using a natural generalization of the RMSD, which we term OT-RMSD:

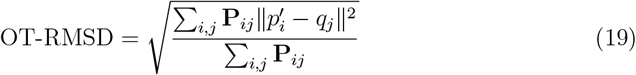

Where 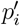 are the coordinates of the mobile structure after the current transformation. This metric represents the expected RMSD under the soft assignment **P**.

After convergence, we extract a final, unambiguous one-to-one alignment (a “hard” correspondence) from the final transport plan **P**. This is achieved via a greedy decoding algorithm that selects the most confident pairings in a one-to-one fashion. The standard RMSD is then calculated on this final “core” set of aligned atoms, providing a single, interpretable metric of structural similarity.

## 3 Results

### 3.1 Experimental Setup

To evaluate the performance of our OT-RMSD method, we conducted a comparative analysis on a benchmark dataset of 40 protein pairs. This set, derived from the RPIC database [2], includes pairs with varying degrees of structural similarity and length differences, providing a challenging test for alignment algorithms.

We compared the performance of three distinct methods:

1. **OT-RMSD (Our Method)**: The algorithm described in this paper, utilizing the simple, distance-based cost function. This corresponds to the ‘OT-simple’ results in our analysis.
2. **OT-heuristic**: For comparison, we also tested a variant of our method that uses a more complex, heuristic-based reward function inspired by the scoring in TM-align. The reward is defined as 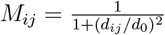, where *d*_*ij*_ is the distance between atoms *i* and *j*, and *d*_0_ is a distance cutoff dependent on the protein length. This method is designed to evaluate the performance of a more traditional, heuristic-based approach within our OT framework.
3. **PyMOL (Baseline)**: A widely-used molecular visualization and analysis tool. We used its standard ‘align’ command with C-alpha atoms, which performs iterative outlier rejection, as a baseline representing a standard, publicly available method.

For each of the 40 pairs, we ran all three algorithms and recorded the number of C-alpha atoms in the final core alignment and the corresponding final RMSD.

### 3.2 Quantitative Analysis of OT-RMSD

The complete results of our benchmark comparison are presented in Table 1. The table provides a side-by-side comparison of the number of aligned atoms and the final RMSD for each of the three methods across all 40 protein pairs. The final row of the table summarizes the performance by showing the arithmetic mean for each metric.

**Table 1:**
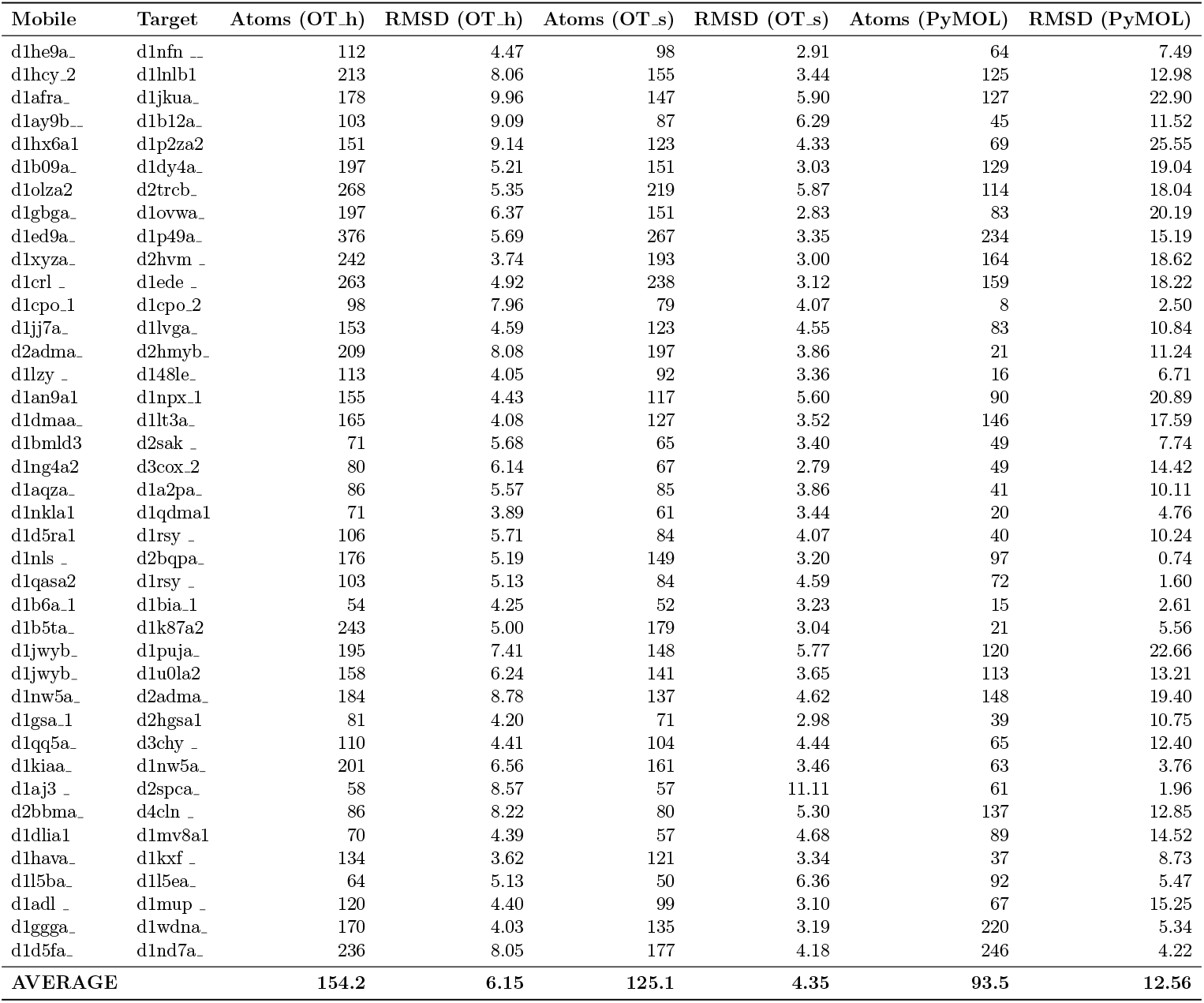
Comparative analysis of alignment algorithms on 40 protein pairs. “Atoms” refers to the number of C-alpha atoms in the final core alignment. “RMSD” is the final root-mean-square deviation in Angstroms (Å) on that core. OT _h is the OT-heuristic variant; OT s is our proposed OT-RMSD method.

An analysis of the average performance, shown in the final row of Table 1, reveals a clear trend. Our proposed OT-RMSD method achieves an average core RMSD of **4.35 °A** across the 40 pairs. This is a marked improvement over the OT-heuristic variant (6.15 °A) and a dramatic improvement over the PyMOL baseline (12.56 °A). In terms of alignment size, OT-RMSD identifies an average core of **125.1 atoms**. This is substantially larger than the PyMOL baseline (93.5 atoms), indicating that OT-RMSD not only produces more accurate alignments but also more comprehensive ones.

### 3.3 Comparison with a Heuristic-Based Cost Function

A key claim of this paper is that a simple, distance-based cost function is superior to more complex, heuristic-driven ones within the OT framework. The comparison between the “OT-RMSD” and “OT-heuristic” columns in Table 1 provides strong evidence for this claim. While the OT-heuristic method, which incorporated ideas from the popular TM-score, was designed with the intention of being a “smarter” algorithm. However, our results show that these additional heuristics, while leading to larger alignments, ultimately degraded the geometric accuracy of the core alignment. This suggests that for the goal of finding a low-RMSD core, a pure, unadulterated geometric cost function is the most effective guide.

### 3.4 Comparison with PyMOL Baseline

When compared to the industry-standard PyMOL ‘align’ command, the advantages of the OT-RMSD framework are even more pronounced. On average, OT-RMSD achieves an RMSD that is nearly three times lower than PyMOL’s (4.35 °A vs 12.56 °A), while simultaneously aligning approximately 34

## 4 Discussion

The results presented in this study highlight the power and elegance of using a principled, first-principles approach to tackle the long-standing problem of protein structure alignment. Our central finding is that a simple, distance-based cost function, when embedded within a robust Optimal Transport framework, not only generalizes the concept of RMSD but also outperforms more complex methods in identifying high-accuracy structural alignments.

The core philosophy of our OT-RMSD method is the “power of simplicity.” By resisting the temptation to build a complex, heuristic-laden scoring function, we allow the mathematics of Optimal Transport to find the most economical mapping between two structures based purely on their geometry. The superior performance of this simple method over the OT-heuristic variant is particularly revealing. The OT-heuristic method, which incorporated ideas from the popular TM-score, was designed with the intention of being a “smarter” algorithm. However, our results show that these additional heuristics, while leading to larger alignments, ultimately degraded the geometric accuracy of the core alignment. This suggests that for the goal of finding a low-RMSD core, a pure, unadulterated geometric cost function is the most effective guide.

The comparison with PyMOL’s ‘align’ command further validates our approach. While PyMOL is an indispensable tool for structural biologists, its default alignment algorithm appears to be tuned for a different objective, often producing larger but less precise alignments. OT-RMSD consistently finds a more geometrically coherent core alignment, as evidenced by its significantly lower average RMSD on a comparable or larger set of atoms. This makes OT-RMSD a powerful complementary tool for researchers seeking to quantify the precise geometric similarity between the cores of two proteins.

Of course, our work is not without limitations. The current implementation is focused on C-alpha atoms and does not account for side-chain orientations. The choice of the sharpness parameter, *γ*, can influence the results, and a more systematic exploration of its effects is warranted. Furthermore, the performance on extremely large protein complexes has not been extensively tested.

Future work could proceed in several exciting directions. Extending the framework to all-atom models is a natural next step. Exploring different ground costs within the OT framework, such as those that incorporate local geometry or flexibility, could yield algorithms tailored for different biological questions. Finally, developing a more robust method for decoding the final hard assignment from the transport plan could further improve the accuracy and coverage of the final alignment.

## 5 Conclusion

In this paper, we have introduced OT-RMSD, a novel method that successfully generalizes the classic Root-Mean-Square Deviation and the Kabsch algorithm to handle the alignment of protein structures of unequal length and unknown correspondence. By leveraging the mathematical framework of Optimal Transport with a simple and fundamental distance-based cost function, our method unifies the search for correspondence and optimal superposition into a single, iterative process. Our benchmark results demonstrate that this principled and parameter-light approach is highly effective, consistently identifying larger and more accurate structural cores than both the widely-used PyMOL ‘align’ command and a more complex, heuristic-driven variant. The success of OT-RMSD underscores the power of combining fundamental geometric principles with modern optimization techniques, providing a robust and valuable tool for quantitative analysis in structural biology.

